# Production and composition of group B streptococcal membrane vesicles varies across diverse lineages

**DOI:** 10.1101/2021.08.24.457602

**Authors:** Cole R. McCutcheon, Jennifer A. Gaddy, David M. Aronoff, Margaret G. Petroff, Shannon D. Manning

## Abstract

Although the neonatal and fetal pathogen Group B *Streptococcus* (GBS) asymptomatically colonizes the vaginal tract of ∼30% of pregnant women, only a fraction of their offspring develops invasive disease. We and others have postulated that these dimorphic clinical phenotypes are driven by strain variability; however, the bacterial factors that promote these divergent clinical phenotypes remain unclear. It was previously shown that GBS produces membrane vesicles (MVs) that contain active virulence factors capable of inducing adverse pregnancy outcomes. Because the relationship between strain variation and vesicle composition or production is unknown, we sought to quantify MV production and examine the protein composition, using label-free proteomics on MVs produced by diverse clinical GBS strains representing three phylogenetically distinct lineages. We found that MV production varied across strains, with certain strains displaying nearly two-fold increases in production relative to others. Hierarchical clustering and principal component analysis of the proteomes revealed that MV composition is lineage-dependent but independent of clinical phenotype. Multiple proteins that contribute to virulence or immunomodulation, including hyaluronidase, C5a peptidase, and sialidases, were differentially abundant in MVs, and were partially responsible for this divergence. Together, these data indicate that production and composition of GBS MVs vary in a strain-dependent manner, suggesting that MVs have lineage-specific functions relating to virulence. Such differences may contribute to variation in clinical phenotypes observed among individuals infected with GBS strains representing distinct lineages.

## Introduction

Group B *Streptococcus* (GBS) is an opportunistic pathogen that asymptomatically colonizes ∼30% of women either vaginally or rectally (1). In individuals with a compromised or altered immune state, including pregnant women, neonates, the elderly, and people living with diabetes mellitus, GBS can cause severe infections (1). Presentation of disease is variable between individuals: in elderly patients and neonates, GBS infection typically presents as septicemia, whereas in pregnant women it more commonly causes chorioamnionitis, preterm birth, or stillbirth (2, 3).

Despite the high prevalence of GBS colonization during pregnancy, only a fraction of babies born to colonized mothers develop an infection. In the United States pregnant individuals colonized with GBS are given antibiotics to reduce the risk of neonatal GBS infection, but even without such prophylaxis most neonates born to GBS-colonized mothers remain infection-free (4). The factors that determine whether a neonate develops GBS sepsis or not are incompletely understood, but evidence implicates bacterial strain variation as a key factor. For example, certain polysaccharide capsular serotypes of GBS are much more common at causing perinatal infections than others (5).

Outside of capsular serotyping, the application of multilocus sequence typing (MLST) has demonstrated that GBS isolates comprise multiple sequence types (STs) that are differentially correlated with disease outcomes (6). While ST-12 strains have been associated with asymptomatic colonization (7), ST-1 and ST-17 strains have been linked to invasive disease in adults and neonates, respectively (6, 8, 9). Moreover, our group has previously shown that different STs interact variably with host cells. ST-17 strains, for instance, had an enhanced ability to attach to gestational tissues, elicited stronger proinflammatory responses, and could persist longer inside macrophages than other STs (10-12). Conversely, ST-12 strains were found to display increased tolerance to ampicillin relative to ST-17 strains (12), highlighting the divergence of these lineages and variation in the ability to withstand different stressors. The mechanisms underlying these strain-dependent differences, however, are poorly understood.

Many bacteria produce membrane vesicles (MVs) of varying sizes (20-500 nm) containing toxins and other virulence factors that can modulate immune responses and influence pathogenesis (13). In addition, GBS can produce MVs that have been implicated in driving infection risk, though this remains an area in need of more research (14, 15). While the exact role of GBS MVs in pathogenesis is not clear, intra-amniotic injection of GBS MVs produced by an invasive ST-7 strain induced preterm birth and intrauterine fetal death in mice (14). GBS MVs were also found to contain active virulence factors that could weaken murine gestational membranes, stimulate immune cell recruitment, and lyse host cells (14, 15). Hence, an important, unanswered question is whether MVs derived from strains belonging to distinct phylogenetic lineages and clinical sources vary in composition and pathogenic potential.

In this study, we sought to compare the quantity and protein composition of MVs produced by genetically distinct GBS strains and evaluate the relationships between proteomic profiles, strain characteristics, and clinical presentation. To accomplish these goals, we isolated MVs from six clinical strains representing three phylogenetic lineages (ST-1, ST-12, and ST-17), and used label-free proteomics to define the protein composition. Using this approach, we report that MV production and composition varies in a strain and ST-dependent manner, highlighting the importance of strain diversity on pathogenic potential.

## Methods

### Bacterial strains

GBS strains GB0020, GB0037, GB0112, GB0411, GB0653, and GB1455 were isolated as described (16, 17); the strain names have been abbreviated for clarity. The invasive isolates GB37, GB411, and GB1455 were isolated from the blood or cerebrospinal fluid of infants with early onset GBS disease (16), while the colonizing isolates GB20, GB112, and GB653 were isolated from vaginal/rectal swabs from asymptomatically colonized mothers before or after childbirth (17). These isolates were previously characterized by MLST and capsular (cps) serotyping (7, 9) and represent the following three common ST, serotype combinations: ST-1, cpsV (GB20, GB37), ST-12, cpsII (GB653, GB1455), and ST-17, cpsIII (GB112, GB411). Strains were cultured using Todd-Hewitt Broth (THB) or Todd-Hewitt Agar (THA) (BD Diagnostics, Franklin Lakes, New Jersey, USA) overnight at 37°C with 5% CO_2_.

### Membrane vesicle (MV) isolation and purification

The isolation and purification of MVs was performed as described (14, 18-20), with some modifications. Briefly, overnight THB cultures were diluted 1:50 into fresh broth and grown to late logarithmic phase (optical density at 600 nm, OD_600_ = 0.9). Aliquots of culture were serially diluted and plated on THA for bacterial enumeration. Cultures were centrifuged at 2000 x g for 20 minutes at 4°C. Supernatants were collected and re-centrifuged at 8500 x g for 15 minutes at 4°C, followed by filtration through a 0.22 µm filter and concentration using Amicon Ultra-15 centrifugal filters (10k Da cutoff) (Millipore Sigma, Burlington, MA, USA). Concentrated supernatants were subjected to ultracentrifugation for 2 hours at 150,000 x g at 4°C. For quantification, pellets were washed by resuspending in PBS, re-pelleting at 150,000 x g at 4°C, and resuspending in PBS; pellets were stored at -80°C until usage.

For proteomics, pellets were resuspended in PBS and purified using qEV Single size exclusion columns (IZON Science, Christchurch, New Zealand) per the manufacturer’s instructions. MV fractions were collected and re-concentrated using the Amicon Ultra-4 centrifugal filters (10 kDa cutoff) (MilliporeSigma, Burlington, Massachusetts, USA) and brought to a final volume of 100 μL in PBS. To preserve the integrity of vesicle proteins, ProBlock Gold Bacterial Protease Inhibitor Cocktail (GoldBio, St. Louis, Missouri, USA) was added. MVs were stored at -80°C until usage.

### Electron microscopy

To visualize GBS MVs, scanning electron microscopy (SEM) was performed on bacterial cultures grown to stationary phase in THB. Culture aliquots were fixed in equal volumes of 4% glutaraldehyde in 0.1 M phosphate buffered saline (pH 7.4), placed on poly-L-lysine coated 12 mm coverslips, and incubated for 5 minutes. The coverslips were washed with water and dehydrated through increasing concentrations of ethanol (25%, 50%, 75%, 95%) for five minutes in each followed by three 5-minute changes in 100% ethanol. Samples were dried in a Leica Microsystems (model EM CPD300) critical point drier using liquid carbon dioxide as the transitional field. Lastly, samples were mounted on aluminum stubs using epoxy glue (System Three Quick Cure 5, System Three Resins, Inc, Lacey, Washington, USA) and coated with osmium (∼10 mm thickness) using a NEOC-AT osmium coater (Meiwafosis Co., Ltd, Tokyo, Japan). Imaging was performed using a JEOL 7500F scanning electron microscope.

To evaluate MV morphology and purity, transmission electron microscopy (TEM) was performed on purified vesicles as described (19). MVs were fixed in 4% paraformaldehyde, loaded onto formvar-carbon coated grids, and counterstained with 2.5% glutaraldehyde and 0.1% uranyl acetate in PBS. Samples were imaged using a JEOL 1400 Flash transmission electron microscope.

### Quantification of vesicle production

Nanoparticle tracking analysis was performed to quantify MVs produced by each strain (n=8-9 replicates per strain) using a NanoSight NS300 (Malvern Panalytical Westborough, MA, USA) equipped with an automated syringe sampler as described previously (19, 21). For each sample, MVs were diluted in phosphate buffered saline (1:100 – 1:1000) and injected with a flow rate of 50. Once loaded, five 20-second videos were recorded at a screen gain of 1 and camera level of 13. After capture, videos were analyzed at a screen gain of 10 and a detection threshold of 4 and data were subsequently exported to a CSV file for analysis using the R package tidyNano (accessed via: https://nguyens7.github.io/tidyNano) (21). Total MV counts were normalized by dividing by the colony forming units (CFUs) of each original bacterial culture.

### Proteomics

Proteomic LC-MS/MS analysis of MVs was performed in duplicate or triplicate by the Proteomics Core at the Michigan State University Research Technology Support Facility (RTSF). Protein concentrations of purified MVs were determined using the Pierce Bicinchoninic Acid Assay (ThermoFisher Scientific, Waltham, Massachusetts) supplemented with 2% SDS in water to reduce the background signal from excess lipids contained within the vesicles. MVs (1.5 μg) were concentrated into a single band in a 4-20% Tris-Glycine SDS-PAGE gel (BioRad, Hercules, CA) that was fixed and stained using colloidal Coomassie blue (22).

Protein bands were excised from the gels and stored in 5% acetic acid at 4°C. Prior to analysis, in-gel trypsin digest and peptide extraction were performed. Briefly, gel bands were dehydrated twice using 100% acetonitrile and incubated with 10 mM dithiothreitol in 100 mM ammonium bicarbonate (pH∼8.0) at 56°C for 45 minutes. Bands were incubated in the dark with 50 mM iodoacetamide in 100 mM ammonium bicarbonate for 20 minutes followed by another dehydration. Sequencing grade modified trypsin (0.01 μg/uL in 50 mM ammonium bicarbonate) was added to each gel band and incubated at 37°C overnight. Peptides extracted by bath sonication (in 60% acetonitrile, 1% trichloroacetic acid solution) were vacuum dried and resuspended (in 2% acetonitrile/0.1% trifluoroacetic) prior to separation using a Thermo ACCLAIM C18 trapping column. Peptides were sprayed onto a ThermoFisher Q-Exactive HFX mass spectrometer for 90 minutes; the top 30 ions per survey were analyzed further using high energy induced dissociation. MS/MS spectra were converted into peak lists using Mascot Distiller v2.7.0 and searched against a SwissProt database containing all GBS sequences available through the National Center for Biotechnology Information (NCBI; accessed 2/08/2019). Contaminants were identified using Mascot searching algorithm v2.7.0, while protein identities were validated using Scaffold v4.11.1.

### Data analysis

To compare MV proteins between strains, proteomic data from all strains were compiled and normalized for inter-experimental variability using Scaffold. Only proteins with a minimum of two identified peptides falling above a 1% false discovery rate and 95% protein threshold, were considered for downstream analysis. Proteins identified as contaminants (via the Mascot searching algorithm v 2.6.0) were removed, whereas proteins identified in both replicates for at least one strain were classified as MV-associated. Subcellular localization analysis was performed using pSORTdb (https://db.psort.org) with protein localization data for GBS strain 2603VR (downloaded from pSORTdb on 3/6/2021). Data visualization and statistical analyses were performed using R version 4.1.0 (https://www.R-project.org). Principle component analysis (PCA) was performed and visualized using the prcomp and fviz_pca functions, respectively. Hierarchical clustering was performed using the pheatmap function and clustered using Euclidean distances. Shapiro tests were used to determine whether data followed a normal distribution and Student t-test (two-sided) or Kruskal-Wallis one-way analysis of variance (ANOVA), in combination with the Dunn’s *posthoc* test, were utilized to test for differences between groups. Multiple hypothesis testing was corrected using Benjamini-Hochberg or Bonferroni correction when necessary.

## RESULTS

### MV production varies across GBS strains

Visualization using SEM revealed abundant production of MVs by all six strains; these MVs were closely associated with bacterial cells (**Figure 1A-B, Figure S1**). Within a given culture, however, some cells displayed a relatively greater number of MVs on the cell surface (**Figure S1**). While rare, these “hyper-producers” were observed in different samples and strains. In addition, TEM revealed that MVs displayed a spherical morphology containing a lipid bilayer and slightly electron dense interior (**Figure 1C-D, Figure S2**), which is typical of bacterial-derived MVs (14, 15).

**Figure 1:**
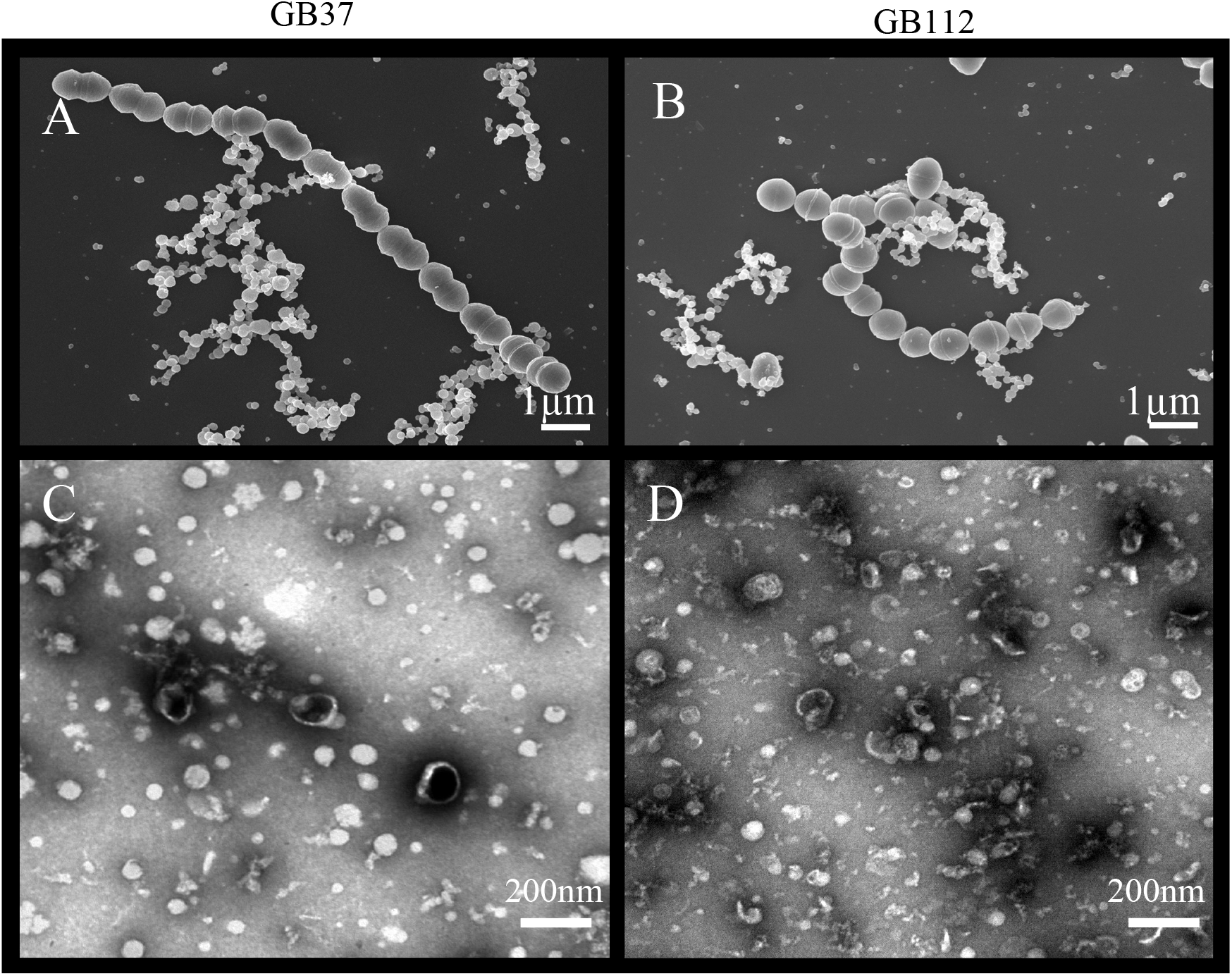
Electron microscopy of membrane vesicles (MVs) from overnight cultures post-purification. Overnight cultures of GBS strains were visualized by electron microscopy. Representative images include the: **A**) invasive ST-1, cpsV (GB37) strain, and **B**) colonizing ST-17, cpsIII (GB112) strain examined by scanning electron microscopy (SEM) at 10,000x magnification with a minimum of 2 replicates per strain. SEM scale bars indicate 1 μm length. Representative transmission electron microscopy (TEM) images of MVs from the same **C**) invasive and **D**) colonizing strains following purification using ultracentrifugation and size exclusion chromatography (2-3 replicates per strain). TEM images were taken at a magnification of 20,000x and the scale bars indicate a length of 200 nm.

Because electron microscopy suggested differences in MV production across strains, we used NanoSight analysis to quantify MV size and production. MVs from each of the six strains displayed a uniform size distribution, ranging between 100 and 200 nm (**Figure 2A**). Similar size distributions were also observed by ST. For MV quantification, total MV counts were normalized to the number of CFUs in the original bacterial cultures. Among the six strains, the average number of MVs/CFU was 0.108 with a range of 0.048-0.206 MVs/CFU; however, considerable variation was detected between strains (**Figure 2B**). Although no difference in MV quantity was observed in colonizing versus invasive strains belonging to ST-1 or ST-17, the ST-1 strains produced significantly fewer MVs relative to the ST-17 strains (**Figure S3**; p <0.0001). While the colonizing ST-12 (cpsII) GB653 strain produced similar vesicle quantities as the two ST-17 (cpsIII) strains, the invasive ST-12 (cpsII) isolate, GB1455, produced significantly more MVs than all other strains examined (p <0.05). By contrast, the colonizing ST-1 (cpsV) isolate, GB20, produced significantly fewer MVs compared to the strains representing all other STs (p <0.05).

**Figure 2:**
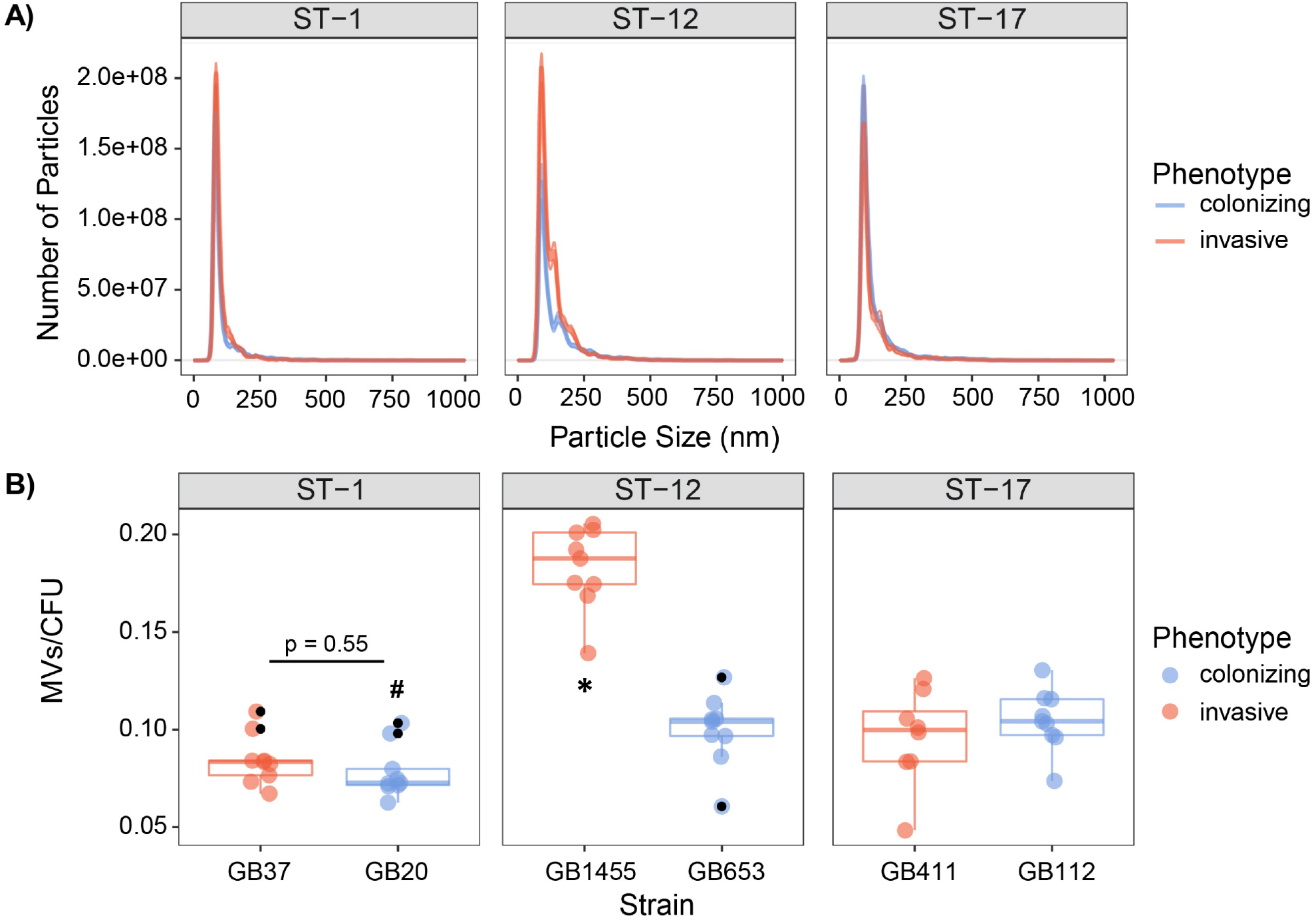
Quantitative assessment of membrane vesicle (MV) production across strains. MVs were isolated by differential centrifugation and quantified using NanoSight analysis. The vesicle **A**) size distribution and **B**) number per bacterial colony forming units (CFUs) are shown for the invasive and colonizing strains by sequence type (ST). For panel B, the lines show the mean across 8-9 biological replicates (indicated by colored dots). Shaded regions surrounding the lines are the standard error of the mean and the black dots are outliers identified by multiplying the interquartile range by 1.5, which was used to extend the upper and lower quartiles. Outliers were excluded from the analysis. Differences in production were assessed using the Kruskal Wallis test followed by a *posthoc* Dunn’s Test with a Benjamini-Hochberg correction. *p-value <0.05 with higher production for all possible comparisons unless otherwise indicated, while # indicates a p-value <0.05 with lower production.

### The MV proteome differs across GBS strains

Proteomics of purified MVs identified 643 total proteins among the six isolates with an average of 458 proteins per strain and range of 239-614 proteins per strain (**Table S1A**). Of note, the number of unique proteins varied by strain. MVs from ST-1 strains, for instance, had fewer unique proteins relative to the other STs with an average of 281 proteins compared to 601 and 493 for the ST-12 and ST-17 strains, respectively. Regardless of ST, however, pSORTdb predicted numerous proteins to be membrane (12-17%) and cell wall (2-11%) localized, while 22-52% were predicted to be localized in the cytoplasm (**Figure 3A**). Although many proteins had a predicted subcellular localization, a large proportion of proteins had unidentified or unpredicted subcellular localization.

**Figure 3.**
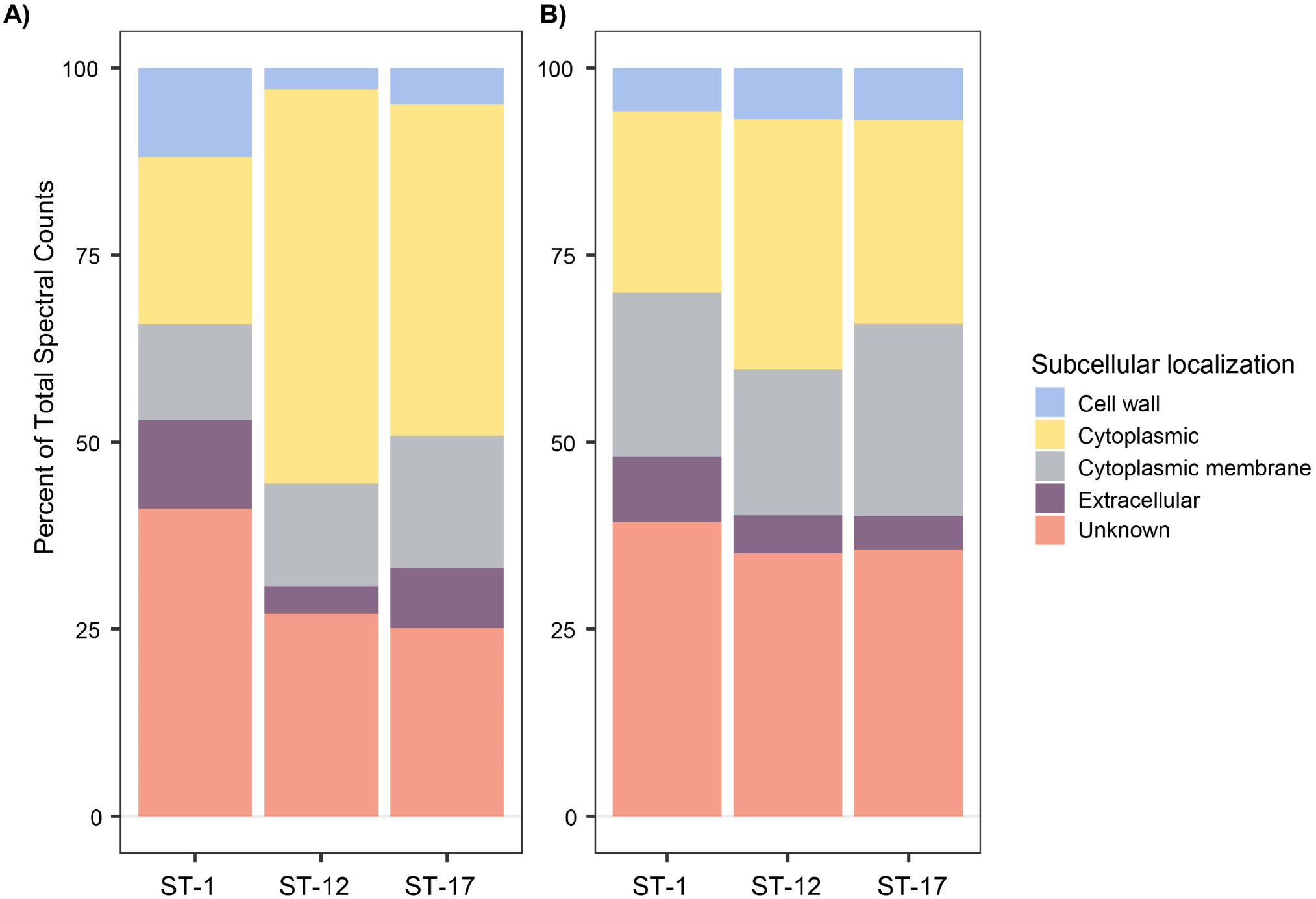
Subcellular localization analysis of membrane vesicle (MV) proteomes. The subcellular localization of **A**) all 643 MV proteins identified, and **B**) a subset of 62 shared MV proteins identified using a pSORTdb database for published *Streptococcus agalactiae* sequences (accessed 3/3/21). Percentages were determined from mean spectral counts for a given sequence type (ST).

Among the total proteins detected, 62 were found in all biological replicates for the six strains (**Table S1B**). These proteins did not vary in spectral abundance between STs and therefore represent the shared MV proteome. Of these 62 proteins, 11 were highly abundant with a mean spectral count greater than 50 (**Table S1C**). Putative, uncharacterized transporters constituted many of these shared proteins, accounting for 39-44% of membrane protein spectral counts. In addition, 19-25% of spectral counts were predicted to have a membrane associated subcellular localization (**Figure 3B**).

Next, we examined whether these proteins were strain-specific or if they were shared in the six strains examined. Of all 643 proteins detected, 192 (29.9%) were detected in at least one biological replicate for all six strains regardless of the clinical phenotype or ST (**Figure 4**). In addition, 124 (19.28%) proteins were shared by the four ST-12 and ST-17 strains but were absent in the ST-1 strains, suggesting that the ST-1 MVs have a unique protein composition. While a minor proportion of proteins were ST- or strain specific, none were shared by all invasive or all colonizing strains.

**Figure 4.**
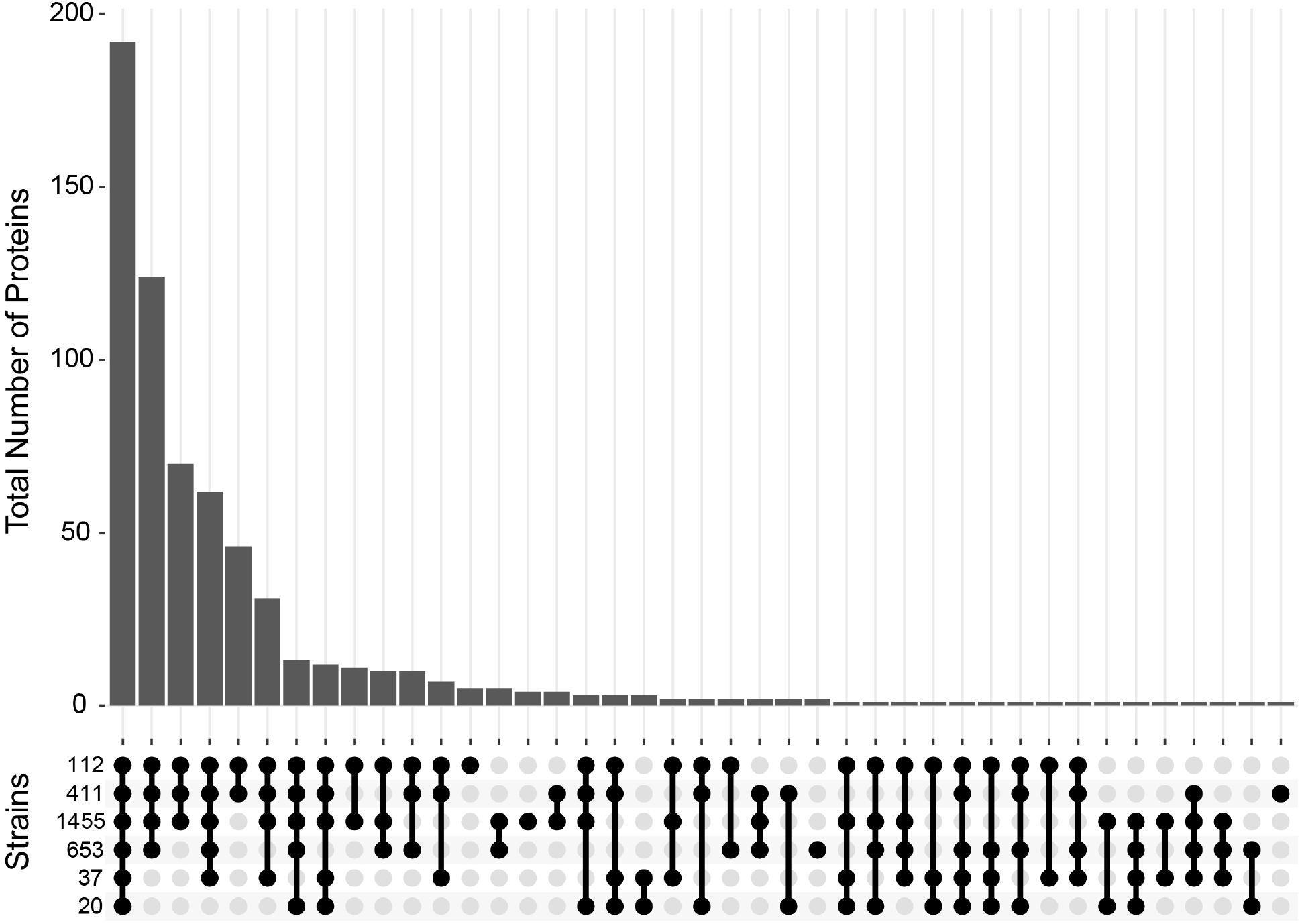
Distribution of proteins detected in membrane vesicles (MVs) among six strains. An Upset plot was generated to show the distribution of all 643 proteins detected across the six GBS strains examined. The y-axis indicates the total number of proteins detected for a given set of strains. Protein presence is defined as having a non-zero spectral count for a given protein in at least one biological replicate for a specific strain. The matrix at the base of the plot shows the strains ordered vertically by sequence type with filled bubbles indicating which strains are positive for the number of proteins detected, and overlaid bars representing number of shared proteins.

We next considered the relationship between protein composition and strain characteristics using PCA. Even though the protein composition of MVs from invasive and colonizing strains overlapped, it was segregated by ST (**Figure 5**), though some overlap was observed between the ST-12 confidence ellipse and those for other STs. No overlap, however, was seen between the ST-1 and ST-17 strains, highlighting their distinct proteomes. This distinct clustering was not observed when the relationship between protein composition and clinical phenotype was analyzed (**Figure S4**), with invasive and colonizing samples displaying a high degree of overlap with little to no separation of their respective confidence ellipses.

**Figure 5:**
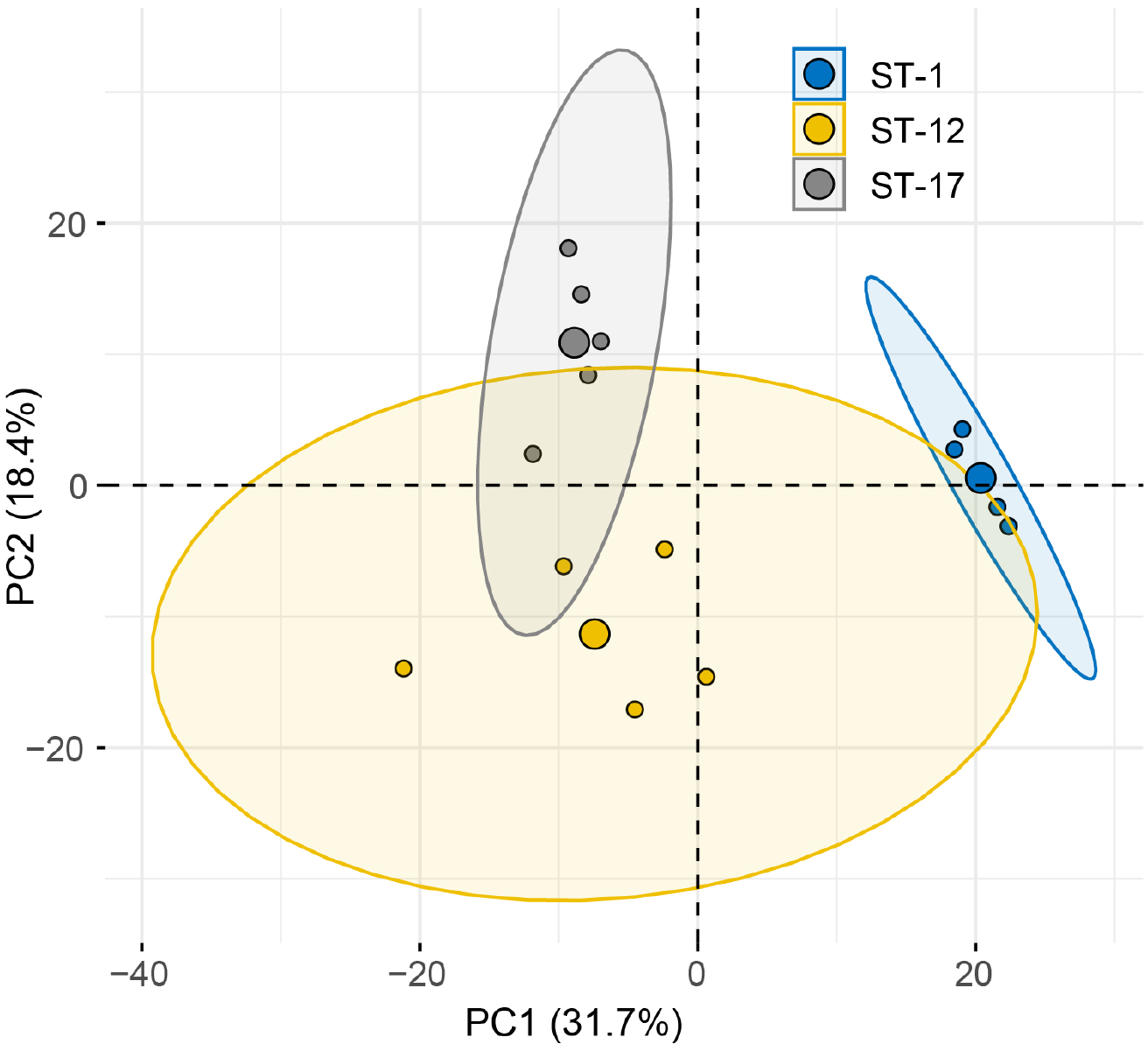
Principal component analysis (PCA) reveals lineage-specific clustering of membrane vesicle (MV) proteomes. PCA of the MV proteomes produced by six strains stratified by sequence type (ST). The large central dot of each ellipse represents the mean point of the corresponding 95% confidence ellipse, while the smaller points represent individual proteomic samples. Confidence ellipses comprise 95% of the samples based on the underlying distribution. Axes percentages represent the amount of variation accounted for by each principal component (PC).

Hierarchical clustering of the protein data further demonstrated that MVs from strains belonging to the same ST had similar protein profiles forming distinct clusters by ST regardless of the clinical phenotype (**Figure 6**). Specifically, proteins from the ST-12 and ST-17 strains formed a distinct branch in the phylogeny that was separate from the ST-1 proteins, indicating that their protein composition was more similar to each other than to ST-1 strains. This observation supports the PCA, showing a higher degree of overlap between ST-12 and ST-17 strains compared to ST-1 strains. Nonetheless, ST-12 and ST-17 strains were still distinguishable, with distinct nodes forming based on protein composition, indicating their divergent composition. This analysis further revealed that ST-1 strains lacked several proteins that were highly abundant in both the ST-12 and ST-17 strains. To a lesser degree than ST-1 MVs, we observed that several highly abundant proteins found among the ST-17 strains were entirely absent in ST-12 strains.

**Figure 6:**
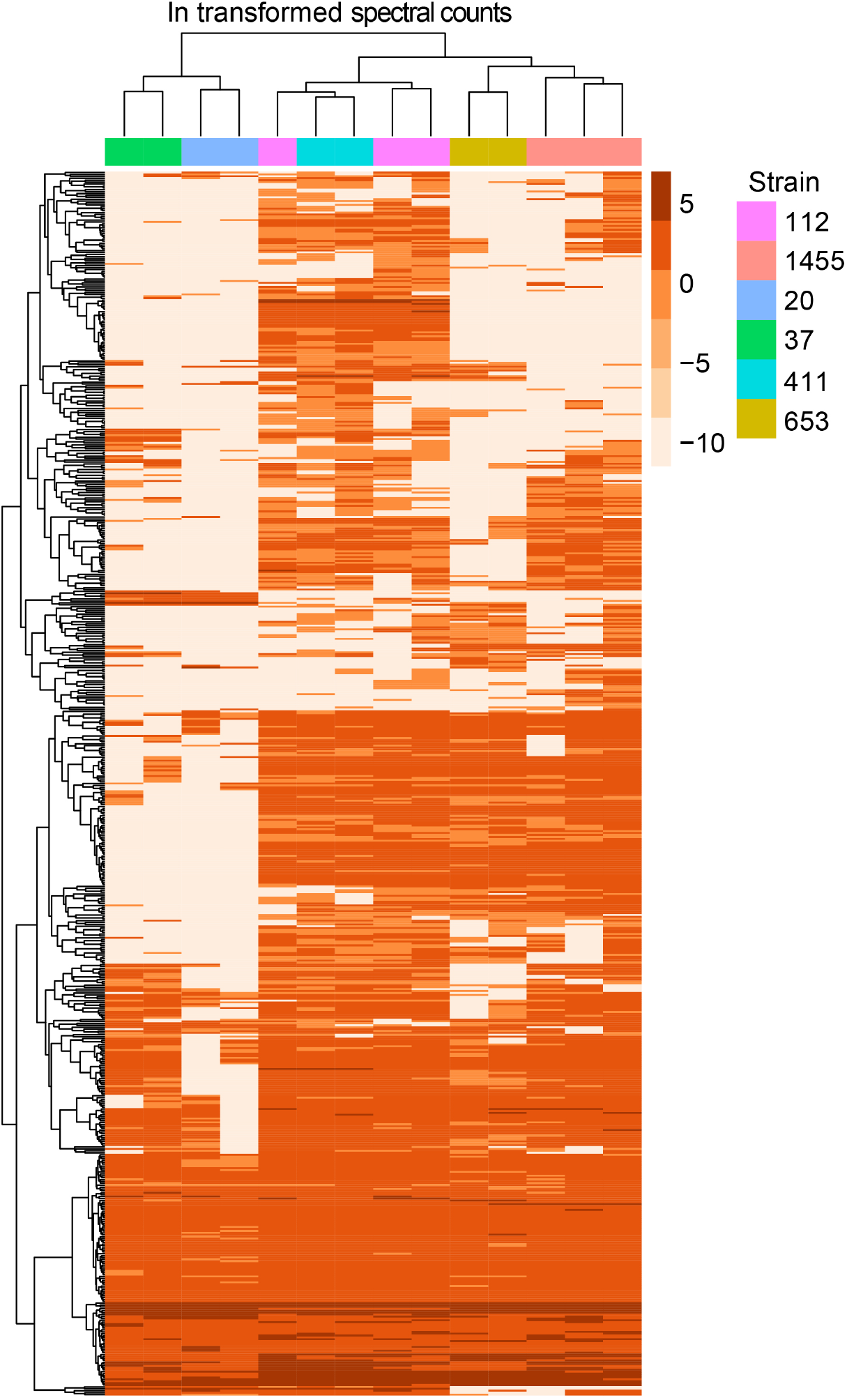
Hierarchical clustering of membrane vesicle (MV) proteomes shows ST specific clustering. A heatmap was generated using hierarchical clustering with the pheatmap function in R, which uses Euclidean distance to cluster rows and columns with similar profiles. Individual rows represent a single accession number for an identified protein, with the color gradient of individual boxes corresponding to the natural log (Ln) transformation of spectral counts for a given protein of interest. Columns represent a single proteomic sample, which are color coded by strain.

### Key virulence factors were differentially abundant in MVs across GBS lineages

To determine which proteins contributed most to the segregation observed in the PCA and hierarchical clustering analyses, we more thoroughly examined the 335 proteins that were significantly enriched in at least one ST (**Table S2**). Notably, several purported virulence factors including the C5a peptidase, hyaluronidase, and sialidase were highly enriched in a ST-dependent manner (**Figure 7**). Both the hyaluronidase and C5a peptidase were significantly more abundant in the two ST-17 strains compared to the ST-1 and ST-12 strains, whereas the sialidase was detected at significantly higher levels in ST-1 versus ST-12 strains. Several proteins of unknown function were also among the most highly abundant and differentially enriched proteins detected. One hypothetical protein, for instance, was significantly more abundant in the ST-1 strains relative to strains representing the other two lineages (**Figure 7**). Similarly, another hypothetical protein was more abundant in the ST-12 strains (**Figure S5**); however, considerable variation was observed across replicates. Numerous phage-associated proteins including a holin and capsid protein, were also detected and found to be more abundant in the ST-17 strains along with several proteins associated with cell division (**Figure S6)**. For example, the average abundance of cell division proteins FtsE, FtsQ, FtsZ,and FtsY, was significantly greater in the two ST-17 strains compared to those from other lineages. Differences in proteins linked to cell wall modification such as penicillin-binding proteins and capsule biosynthesis proteins, were also detected (**Figure S7)**.

**Figure 7:**
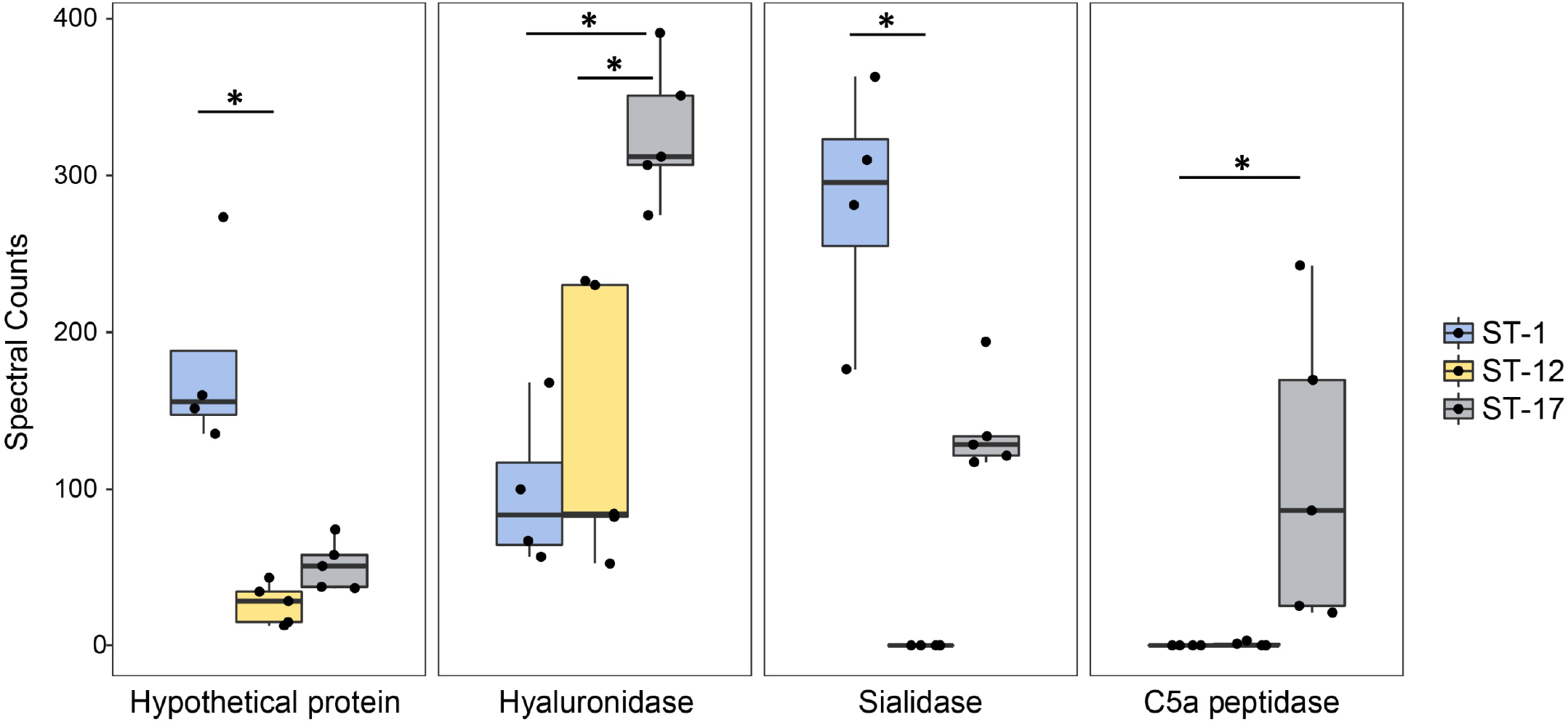
Highly abundant proteins are present at variable levels in membrane vesicles (MVs). The spectral counts of specific proteins were plotted after stratifying by the sequence type (ST). The median spectral count associated with each ST is represented within each box. The black dots represent a single biological replicate for a given strain. Statistical comparison was performed using a Kruskal Wallis test. Multiple pairwise comparisons were then made using the pairw.kw function in R, which uses a conservative Bonferroni correction method to correct for multiple hypothesis testing. Comparisons with p-values < 0.05 are denoted with an asterisk.

## DISCUSSION

Current knowledge regarding GBS derived MVs is restricted to one clinical strain (14, 15) and hence, we sought to examine MV production and composition in a set of clinical strains with different traits. While no clear association was observed between clinical phenotype and the production or composition of MVs, we have demonstrated that the GBS MV proteome is ST-dependent. The same was observed for MV production, though some variation was noted between strains of the same ST. Together, these data indicate that GBS MVs have strain-dependent functions that could impact survival in hosts, immunomodulation, and virulence.

This study expands our current knowledge of GBS MVs by highlighting their potential impact on virulence. Specifically, we demonstrated that GBS MVs have a high abundance of immunomodulatory virulence factors including C5a peptidase, hyaluronidase, and sialidase (23-25). The bifunctional C5a peptidase has been shown to promote the degradation of the proinflammatory complement component (C5a) while simultaneously promoting bacterial invasion into host cells (23, 24). MVs from both ST-17 (cpsIII) strains examined herein contained high levels of C5a peptidase, whereas ST-1 and ST-12 strains lacked this protein. Intriguingly, ST-17 strains were previously shown to possess distinct virulence gene profiles as well as unique alleles of *scpB* encoding the C5a peptidase (26, 27), suggesting that ST-17 strains may be primed to cause invasive infections. This suggestion is in line with epidemiological data showing that ST-17 strains are important for invasive disease in adults and neonates (6, 8, 9) as well as mechanistic studies showing an enhanced ability to attach to gestational tissues, induce stronger proinflammatory responses, and persist inside macrophages (10-12). Nonetheless, it is important to note that our clinical definitions of “invasive” versus “colonizing” strain types may not be representative of each strain population. Although strains isolated from an active infection clearly demonstrate “invasive” potential, it is possible that strains designated as “colonizing” could also cause an infection in specific circumstances and host environments.

Although sialidases have no known role in GBS pathogenesis (25), these proteins were shown to be immunomodulatory in other bacterial species (28, 29) while simultaneously promoting biofilm production and metabolism of host sugars (30, 31). The presence and abundance of sialidase was variable: the ST-1 and ST-17 MVs all contained sialidase, but the ST-12 MVs lacked it. In two prior studies examining GBS MVs produced by a ST-7 strain, A909, neither C5a peptidase nor sialidase were identified (14, 15), further highlighting differences across strains. However, we cannot rule out the possibility that the abundance of these virulence factors was beneath the detection limit in those studies. Similarly, the previous analysis of GBS MVs highlighted the importance of hyaluronidase (14). This immunomodulatory factor has previously been shown to promote ascending infection, degrade host extracellular matrix components, and dampen the host immune response (24). While we also found high levels of hyaluronidase in the ST-17 MVs examined, our results further show that the ST-12 and ST-1 MVs contained significantly lower amounts of this protein.

It is also important to note that multiple uncharacterized and hypothetical proteins were detected. Previous reports have demonstrated that in gram positive species, roughly 30-60% of all MV proteins map to the cytoplasm (32, 33). While our results are consistent with this observation showing ∼22-52% of all proteins mapping to the cytoplasm, roughly 25-41% of the GBS MV proteins had an unidentifiable subcellular localization. Similar trends of ST-dependent enrichment of several hypothetical proteins were observed, with these representing some of the most highly abundant proteins. Although some uncharacterized proteins, such as those classified as putative ABC transporters, have predicted functions, their role in vesicle function or virulence is currently unknown. Future analyses must be undertaken to identify which proteins play a role in MV associated pathogenesis.

Through this study, we have also identified a shared proteome among MVs from phylogenetically distinct GBS strains. In total, 62 proteins were consistently found within GBS MVs regardless of the ST. Indeed, over 17% of these shared proteins were highly abundant, indicating that they may be important for MV functionality. Even though many of these proteins have yet to be characterized, we identified an abundance of transporter proteins in MVs suggesting a potential role in MV function. Separate of functionality, these shared proteins may be of value as potential MV markers in future studies.

While various mechanisms have been proposed for the biogenesis of gram positive MVs, those mechanisms important for GBS MV biogenesis are unclear (13, 34). Our data demonstrate that diverse GBS strains produce MVs with consistent size distributions, indicating that GBS MV production is ubiquitous. Purported mechanisms of MV biogenesis include phage mediated biogenesis (35, 36), membrane budding during division (37), and cell wall remodeling (13, 38). In line with these mechanisms, our proteomics analysis revealed the presence of phage associated proteins, division septum-associated proteins and cell wall-modifying enzymes. Several of these proteins were also differentially abundant, with some proteins being more highly enriched in certain STs. For instance, phage proteins were enriched in ST-17 strains but were nearly absent in ST-12 and ST-1 strains. Although we observed similar enrichment of cell division proteins in ST-12 and ST-17 strains relative to ST-1, cell wall modifying proteins were most abundant in the ST-17 strains. Taken together, these data indicate that MVs are produced by diverse strains with varying traits; however, the mechanisms for MV biogenesis appear to be strain dependent. Additional studies are needed to test this hypothesis.

Although our study has enhanced our understanding of the proteomic composition of GBS MVs, it has a few limitations. Because strains of each GBS lineage possess the same capsule (cps) type, it is difficult to differentiate between ST versus cps effects. Another concern when dealing with MVs is the presence of non-vesicular contaminants. In some eukaryotic and prokaryotic systems where the composition of MVs is well defined, markers are used to assess purity (39-41). Due to the relatively unknown composition of GBS MVs, however, we were unable to target specific markers to evaluate the purity. Rather, we relied on size exclusion chromatography followed by TEM to further remove non-vesicular proteins from each MV preparation. While we likely have some contaminant proteins, the purity of our preparations exceeds those performed in prior GBS studies (14, 15) and mimics protocols optimized for removing extravesicular macromolecules from Gram positive MVs (14, 15, 42, 43). Indeed, studies in *Staphylococcus aureus* and *Streptococcus mutans* have confirmed the presence of similar proportions of cytoplasmic and extracellular proteins within MVs (32, 33). Further, while our study has greatly enhanced our understanding of GBS MV composition, it is known that other macromolecules are present within MVs (14). Whether these macromolecules display ST dependent composition is unclear; however, given these data further studies are warranted.

In summary, this comprehensive analysis of GBS MVs from strains representing three phylogenetically distinct lineages demonstrates strain dependent composition and production of MVs. Our data further demonstrate that MVs carry both known virulence factors and other proteins of unknown function in variable abundance between strains, suggesting that they may have altered functionality or ability to promote virulence. Follow up studies elucidating virulence and immunomodulatory properties of diverse strains of GBS MVs are therefore warranted, particularly given the high level of variation in protein composition observed among only these six strains. Taken together, these findings further highlight the importance of strain variation in GBS pathogenesis and shed light on the potential role of MVs in virulence.

## Acknowledgments

We would like to thank Dr. H. Dele Davies for sharing the bacterial strains and Drs. Sean L. Nguyen and Soo H. Ahn for helpful conversations and assistance with data analysis. We also thank Karla Vasco for assisting with the hierarchical clustering heatmap analysis as well as Alicia Withrow, Carol Flegler, and Douglas Whitten for their assistance with TEM, SEM, and proteomics analysis, respectively.

## Data Availability

Raw proteomic data was submitted to the MassIVE database (massive.ucsd.edu). Data can be accessed via https://doi.org/doi:10.25345/C5RC1H or ftp://massive.ucsd.edu/MSV000087985/.

